# Cell competition between wild-type and JAK2V617F mutant cells in a murine model of a myeloproliferative neoplasm

**DOI:** 10.1101/2020.08.26.267070

**Authors:** Melissa Castiglione, Haotian Zhang, Huichun Zhan

## Abstract

The myeloproliferative neoplasms (MPNs) are clonal stem cell disorders characterized by overproduction of mature blood cells and increased risk of transformation to frank leukemia. The acquired kinase mutation JAK2V617F plays a central role in a majority of these disorders. The hematopoietic stem cell (HSC) compartment in MPN is heterogeneous with the presence of both JAK2 wild-type and JAK2V617F mutant cells in most patients with MPN. Utilizing *in vitro* co-culture assays and *in vivo* competitive transplantation assays, we found that the presence of wild-type cells altered the behavior of co-existing JAK2V617F mutant cells, and a mutant microenvironment (niche) could overcome the competition between wild-type and mutant cells, leading to mutant clonal expansion and overt MPN. We also demonstrated that competition between wild-type and JAK2V617F mutant cells triggered a significant immune response, and there was a dynamic PD-L1 deregulation in the mutant stem/progenitor cells caused by their interactions with the neighboring wild-type cells and the microenvironment. Therefore, while accumulation of oncogenic mutations is unavoidable during aging, our data suggest that, if we could therapeutically enhance normal cells’ ability to compete, we might be better able to control neoplastic cell expansion and prevent the development of a full-blown malignancy.

**Key Points:**

- The presence of wild-type cells alters the behavior of co-existing JAK2V617F mutant cells
- A mutant microenvironment overcomes the competition between wild-type and JAK2V617F mutant cells, leading to the development of a MPN

## Introduction

Cancer is the pathological outcome of competition between wild-type cells and cancerous cells. During their cellular evolution, cells bearing oncogenic mutations develop multiple molecular and cellular mechanisms, both cell intrinsic and cell extrinsic, to make them more competitive than their normal counterparts. Cell competition is an evolutionarily conserved mechanism involved in development, tissue homeostasis, and stem cell maintenance. It is akin to natural selection between species, in that “fitter” cells win out over their “less-fit” neighbors.^1–4^ Such competition is seen only when there is a mixture of genetically different cells, a phenomenon known as mosaicism. Studies from the past several years revealed that human tissues (e.g. skin, esophageal) show high levels of mosaicism with many precancerous mutations, yet such tissues rarely develop frank tumors.^5,6^ These observations suggest that cell competition can protect against cancer.

The myeloproliferative neoplasms (MPNs) are clonal hematopoietic stem/progenitor cell (HSPC) disorders characterized by overproduction of mature blood cells, and increased risk of transformation to acute leukemia. The acquired signaling kinase mutation JAK2V617F plays a central role in most patients with these disorders. The HSPC compartment in MPN is heterogeneous with the presence of both JAK2 wild-type and JAK2V617F mutant cells in most patients with MPN, at least early in their disease process.^7^ Despite mutant cells bearing an *in vitro* proliferative advantage because of constitutive kinase activity, in some patients, there is little or no change in the mutant/wild-type cell ratio over long periods of follow up.^8–10^ In others, the MPN can evolve to acute leukemia and patients experience high relapse rates following allogeneic stem cell transplantation, the only curative treatment for patients with MPNs.^11–14^ JAK2V617F is also one of the common mutations associated with clonal hematopoiesis of indeterminate potential, which is defined as the presence of a somatic mutation in at least 2-4% blood cells without other hematologic abnormalities. Such clonal hematopoiesis is present in 10-20% of people older than 70 yrs and most of these individuals do not convert to advanced disease.^15–19^ These unique features make the JAK2V617F mutant MPN a model disease to study the early stages of clonal competition and tumor progression. In this work, we investigated the competitive interactions between wild-type and JAK2V617F mutant cells, and the impact of microenvironmental factors in driving mutant clonal dominance in the MPNs.

## Methods

### Experimental mice

JAK2V617F Flip-Flop (FF1) mice^20^ was provided by Radek Skoda (University Hospital, Basal, Switzerland) and *Tie2-Cre* mice^21^ by Mark Ginsberg (University of California, San Diego). FF1 mice were crossed with Tie2-Cre mice to express JAK2V617F specifically in hematopoietic cells and endothelial cells (ECs) (Tie2-cre^+^FF1^+^ mice), so as to model the human diseases in which both the hematopoietic stem cells and ECs harbor the mutation.^22–27^ All mice used were crossed onto a C57BL/6 background and bred in a pathogen-free mouse facility at Stony Brook University. CD45.1+ congenic mice (SJL) were purchased from Taconic Inc. (Albany, NY). All mice were fed a standard chow diet. No randomization or blinding was used to allocate experimental groups. Animal experiments were performed in accordance with the guidelines provided by the Institutional Animal Care and Use Committee.

### Complete blood counts and colony assays

Peripheral blood was obtained from the facial vein via submandibular bleeding, collected in an EDTA tube, and analyzed using a Hemavet 950FS (Drew Scientific). Mouse methylcellulose complete media (M3434, Stem Cell Technologies, Vancouver, BC) were used to assay hematopoietic colony formation, which were enumerated according to the manufacturer’s protocol.

### Marrow cell isolation

For marrow cells, murine femurs and tibias were first harvested and cleaned thoroughly. Marrow cells were flushed into PBS with 2% fetal bovine serum using a 25G needle and syringe. Remaining bones were crushed with a mortar and pestle followed by enzymatic digestion with DNase I (25U/ml) and Collagenase D (1mg/ml) at 37 °C for 20 min under gentle rocking. Tissue suspensions were thoroughly homogenized by gentle and repeated mixing using 10ml pipette to facilitate dissociation of cellular aggregates. Resulting cell suspensions were then filtered through a 40uM cell strainer.

For depletion of mature hematopoietic cells, the Lineage Cell Depletion Kit (Miltenyi Biotec, San Diego, CA) was used. The lineage negative cells were collected and then positively selected for CD117^+^ (cKit^+^) cells using CD117 microbead (Miltenyi Biotec) to yield Lin^-^cKit^+^ HSPCs.

### In vitro co-cultures

Marrow CD45^+^CD201^+^CD150^+^CD48^-^ cells, a highly purified long-term repopulating stem cell population,^28,29^ were isolated from wild-type mice (CD45.1) or JAK2V617F-positive Tie2-cre^+^FF1^+^ mice (CD45.2) by flow cytometry. Cells were cultured either directly or in a Transwell unit with 1.0um pore size (Corning, NY) in StemSpan® serum-free expansion medium (SFEM) containing 100 ng/mL recombinant mouse SCF and 100ng/mL recombinant human TPO for a total of 7 days (all from Stem Cell Technologies).

### Stem cell transplantation assays

Recipient mice were irradiated with two doses of 540 rad 3h apart. Donor cells were injected into recipients by standard intravenous tail vein injection using a 27G insulin syringe. For competitive transplantation, 5×10^5^ CD45.2 donor marrow cells from Tie2-cre^+^FF1^+^ mice were injected intravenously together with 5×10^5^ competitor CD45.1 wild-type marrow cells. For non-competitive transplantation, 1×10^6^ unfractionated donor marrow cells were transplanted into wild-type recipients by intravenous tail vein injection. After transplantation, mice were maintained on antibiotic water for 4-6 weeks.

### BrdU incorporation analysis

Mice were injected intraperitoneally with a single dose of 5-bromo-2’-deoxyuridine (BrdU; 100 mg/kg body weight) and maintained on 1mg BrdU/ml drinking water for two days. Mice were then euthanized and marrow cells isolated as described above. For analysis of HSPC proliferation, Lineage^neg^ (Lin^-^) cells were first enriched using the Lineage Cell Depletion Kit (Miltenyi Biotec) before staining with fluorescent antibodies specific for cell surface HSPC markers, followed by fixation and permeabilization using the Cytofix/Cytoperm kit (BD Biosciences, San Jose, CA), DNase digestion (Sigma, St. Louis, MO), and anti-BrdU antibody (Biolegend, San Diego, CA) staining to analyze BrdU incorporation. For analysis of more abundant cell populations, marrow cells were stained with cell surface antibodies, then fixed and stained with anti-BrdU antibody for BrdU incorporation analysis as described above.

### Flow cytometry

All samples were analyzed by flow cytometry using a FACSAria III (BD biosciences, San Jose, CA, USA). CD45 (Clone 104) (Biolegend, San Diego, CA, USA), CD45.1 (Clone A20) (BD Biosciences), CD45.2 (Clone 104) (Biolegend), Lineage cocktail (include CD3, B220, Gr1, CD11b, Ter119, Biolegend), cKit (Clone 2B8, Biolegend), Sca1 (Clone D7, Biolegend), EPCR (CD201) (Clone eBio1560, eBioscience, San Diego, CA, USA), CD150 (Clone mShad150, eBioscience), CD48 (Clone HM48-1, Biolegend), and PD-L1 (Clone 10F.9G2, Biolegend) antibodies were used.

### Cell cycle analysis

For HSPC cell cycle analysis, marrow cells were first stained with fluorescent antibodies for cell surface markers (CD45.1, CD45.2, CD150, and CD48), washed, and then stained with Hoechst33342 (10ug/ml) (Sigma) at 37°C in dark for 45 min, followed by staining with Pyronin Y (0.5ug/ml) (Sigma) at 37°C in dark for another 45 min. Cells were kept on ice until flow cytometry analysis on a LSR II (BD biosciences).^30,31^

### Analysis of apoptosis by active caspase-3 staining

Marrow cells were stained with fluorophore-conjugated antibodies against lineage markers (CD2, CD3, CD5, CD8a, B220, Ter119, and Gr1), cKit, Sca1, CD150, CD48. Cells were then washed and fixed using the BD Fixation/Permeabilization Solution Kit according to the manufacturer’s instruction. Cells were then stained using a rabbit anti-activated caspase-3 antibody. Data were acquired using a FACSAria II flow cytometer.

### Analysis of senescence by senescence associated β-galactosidase (SA-fi-Gal) activity

Marrow cells were stained with fluorophore-conjugated antibodies against lineage markers (CD2, CD3, CD5, CD8a, B220, Ter119, and Gr1), cKit, Sca1, CD150, CD48. Cells were then washed and fixed using 2% paraformaldehyde and incubated with CellEvent™ Senescence Green Probe (ThermoFisher Scientific, Waltham, MA) according to the manufacturer’s instruction. Data were acquired using a FACSAria II flow cytometer.

### Transcriptome analysis of cardiac ECs using RNA sequencing

For RNA sequencing experiments, wild-type and JAK2V617F mutant marrow Lin^-^cKit^+^ HSPCs were isolated by flow sorting and magnetic bead isolation (Miltenyi Biotec, San Diego, CA). Total RNA was extracted using the RNeasy mini kit (Qiagen, Hilden, Germany). RNA integrity and quantitation were assessed using the RNA Nano 6000 Assay Kit of the Bioanalyzer 2100 system (Agilent Technologies, CA, USA). For each sample, 400ng of RNA was used to generate sequencing libraries using NEBNext® Ultra™ RNA Library Prep Kit for Illumina® (New England BioLabs, MA, USA) following manufacturer’s recommendations. The clustering of the index-coded samples was performed on a cBot Cluster Generation System using PE Cluster Kit cBot-HS (Illumina) according to the manufacturer’s instructions. After cluster generation, the library preparations were sequenced on an Illumina platform. Index of the reference genome was built using hisat2 2.1.0 and paired-end clean reads were aligned to the reference genome using HISAT2. HTSeq v0.6.1 was used to count the reads numbers mapped to each gene. Differential expression analysis between pooled wild-type HSPCs (from recipients of wild-type donors, n=3), JAK2V617F mutant HSPCs without competition (from recipients of JAK2V617F mutant donors, n=2), and JAK2V617F mutant HSPCs with competition (from recipients of both wild-type and mutant donors, n=3) was performed using the DESeq R package (1.18.0). The resulting *P*-values were adjusted using the Benjamini and Hochberg’s approach for controlling the false discovery rate. Genes with an adjusted *P*-value < 0.05 found by DESeq were assigned as differentially expressed. Gene Ontology (GO) (http://www.geneontology.org/) enrichment analysis of differentially expressed genes was implemented by the ClusterProfiler R package. GO terms with corrected *P*-value less than 0.05 were considered significantly enriched.

### Statistical analysis

Statistical analysis was performed using Student’s t tests (2 tailed) using Excel software (Microsoft). A p value of less than 0.05 was considered significant. Data are presented as mean ± standard error of the mean (SEM). All experiments were conducted and confirmed in at least two replicates.

## Results

### Competition between wild-type and JAK2V617F mutant HSPCs *in vitro*

To study how wild-type and JAK2V617F mutant cells interact, we isolated murine wild-type and JAK2V617F mutant CD45^+^CD201^+^CD150^+^CD48^-^ cells, a highly purified long-term repopulating stem cell population,^28,29^ from either wild-type mice (CD45.1) or JAK2V617F-positive Tie2-cre^+^FF1^+^ mice (CD45.2)^32–34^ and performed a series of *in vitro* co-culture experiments. We found that JAK2V617F mutant HSPCs displayed a higher proliferation rate than wild-type HSPCs when cultured separately. In contrast, when placed in direct co-culture, proliferation of wild-type HSPCs was greatly increased, such that no significant differences between wild-type and mutant cells was detected. (Figure 1A-B) This was further confirmed by hematopoietic colony formation assays in which cocultured JAK2 wild-type and JAK2V617F mutant HSPCs generated equal numbers of wild-type and mutant colonies. (Figure 1C) Similar results were also obtained when using a transwell co-culture system, (Figure 1D-E) suggesting that direct cell-cell contact was not required for the competitive interactions between the wild-type and mutant HSPCs we have observed in the various co-cultures.

**Figure 1.**
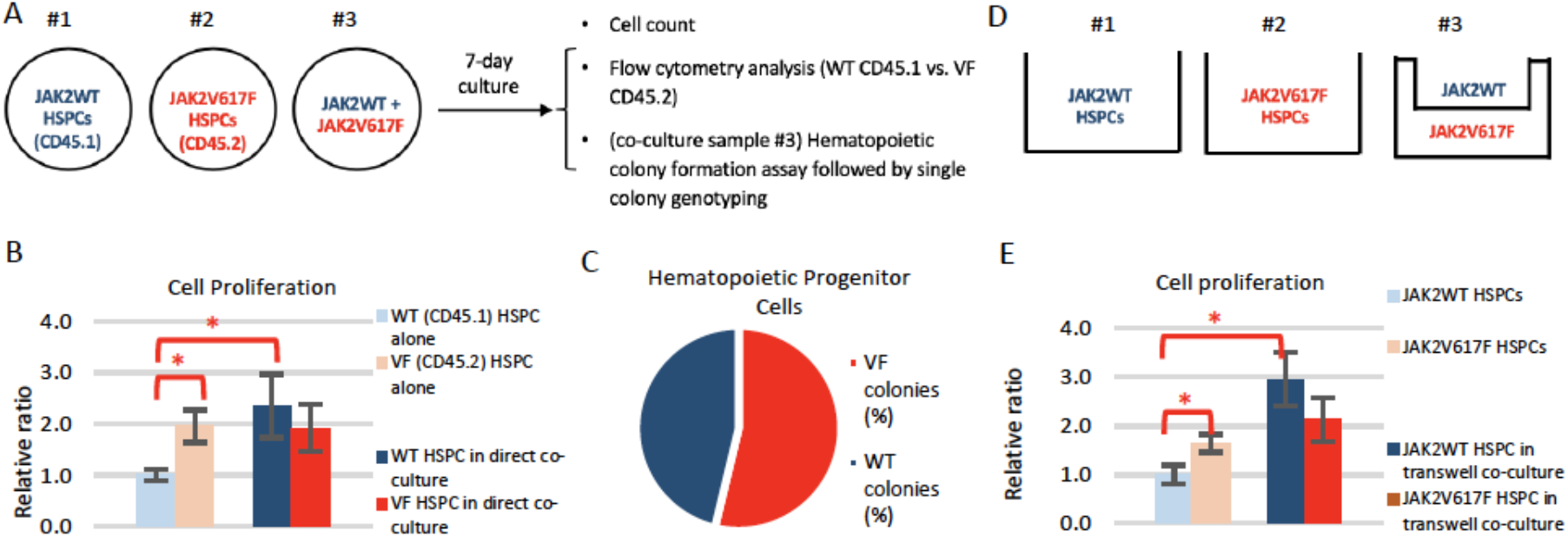
JAK2 wild-type and JAK2V617F mutant cell competition *in vitro.* (**A**) Experimental scheme of direct co-culture. (**B**) Cell proliferation of JAK2 wild-type (WT) and JAK2V617F (VF) HSPCs (CD45^+^CD201^+^CD150^+^CD48^-^) when cultured alone and when cultured together. (**C**) Co-cultured JAK2 wild-type and JAK2V617F HSPCs were plated on methocult for hematopoietic progenitor cell assay. Single colony genotyping demonstrated co-existence of both wild-type and JAK2V617F mutant colonies after co-culture. (**D**) Experimental scheme of transwell co-culture experiment. (**E**) Cell proliferation of JAK2WT and JAK2V617F HSPCs when cultured alone and when cultured together in transwells. \**P*<0.05

### The presence of wild-type cells alters the behavior of co-existing JAK2V617F mutant cells *in vivo*

To investigate the competitions between wild-type and JAK2V617F mutant HSPCs *in vivo,* we performed a series of marrow transplantation experiments. When 100% JAK2V617F mutant marrow cells were transplanted into lethally irradiated wild-type recipients, recipient mice developed a MPN phenotype with leukocytosis and thrombocytosis ~4-8wks after transplantation. In contrast, when 50% mutant marrow cells (CD45.2) and 50% wild-type marrow cells (CD45.1) were transplanted together into lethally irradiated wild-type recipient mice, the JAK2V617F mutant donor cells displayed engraftment similar to the wild-type donor cells, and the recipient mice had normal blood cell counts during more than 4-months of follow up.^32^ (Figure 2A-C) In addition, marrow Lin^-^ cKit^+^Sca1^+^CD150^+^CD48^-^ HSC^35^ frequencies in recipients of 50% mutant and 50% wild-type cells were significantly lower compared to recipients of 100% mutant cells. (Figure 2D) While this appears to be in conflict with other reports that the JAK2V617F-positive MPN phenotype is transplantable, and usually develops as early as 4 weeks following transplantation, in every such study 100% JAK2V617F-positive marrow cells were transplanted into wild-type recipients.^36–40^ These results provide clear evidence that the presence of wild-type cells alters the behavior of co-existing JAK2V617F mutant cells *in vivo.*

**Figure 2.**
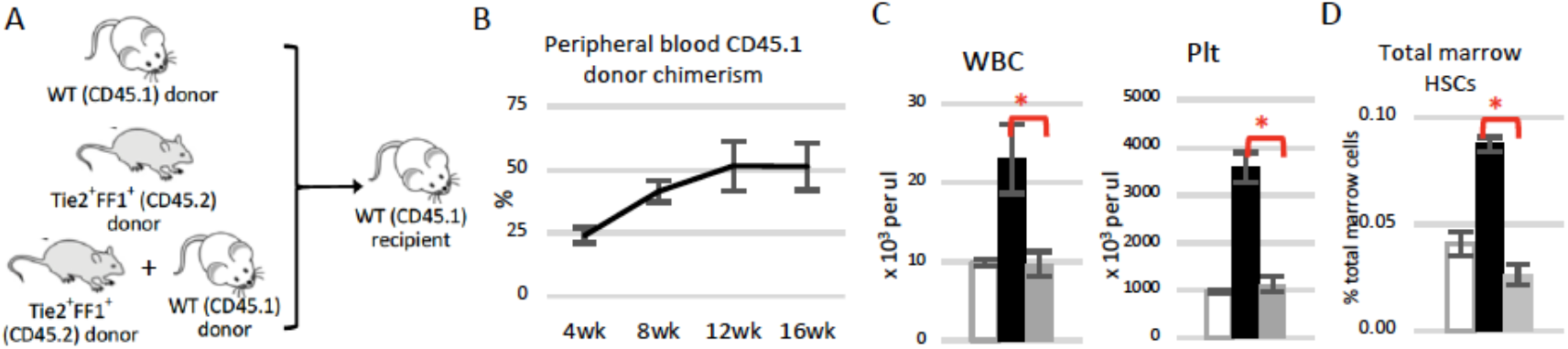
The presence of wild-type cells alters the behavior of JAK2V617F mutant cells *in vivo.* (**A**) Scheme of marrow transplantation experiments. (**B**) Peripheral blood wild-type (CD45.1) donor chimerism after competitive repopulation assay. (**C-D**) Peripheral blood cell counts (C) and marrow Lin^-^cKit^+^Sca1^+^CD150^+^CD48^-^ HSC frequencies (D) in recipient mice of wild-type donor alone (white), JAK2V617F mutant donor alone (black), or both wild-type and JAK2V617F mutant donor cells (grey) 16-20wk after transplantation. (n=5-8 mice in each group in B-C; n=4-5 mice in each group in D). * *P*<0.05

### A mutant microenvironment overcomes the competition between wild-type and JAK2V617F mutant cells, leading to the development of a MPN

Endothelial cells (ECs) are an essential component of the hematopoietic niche and most HSPCs reside close to a marrow sinusoid (the “vascular niche”).^41^ ECs carrying the JAK2V617F mutation can be detected in patients with MPNs.^22,23^ We performed a series of transplantation experiments using the Tie2-cre^+^FF1^+^ murine model which expresses the mutation specifically in all hematopoietic cells (including HSPCs) and vascular ECs.^20,21^ As we previously reported,^32^ when JAK2V617F mutant marrow cells were injected intravenously together with wildtype marrow cells into lethally irradiated Tie2-cre^+^FF1^+^ mice (with mutant ECs) or control mice (with normal ECs), only the Tie2-cre^+^FF1^+^ recipient mice developed a MPN phenotype with mutant HSPC expansion. (Figure 3A-C)

**Figure 3.**
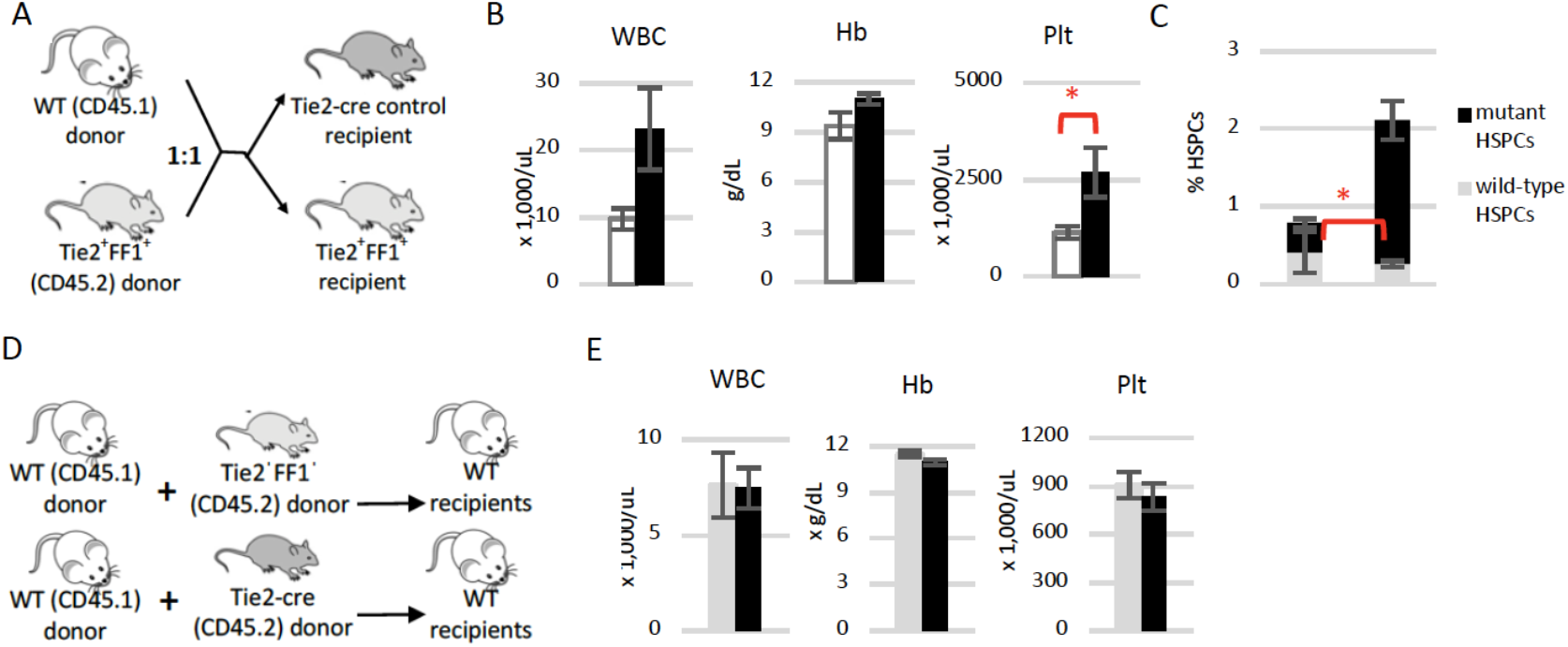
A mutant marrow microenvironment alters the competition between wild-type and JAK2V617F mutant cells. (**A**) Scheme of competitive marrow transplantation experiments where both wild-type and JAK2V617F marrow donors were injected together into lethally irradiated Tie2-cre control mice (with wild-type vascular niche) or Tie2-cre^+^FF1^+^ mice (with JAK2V617F mutant vascular niche). (**B**) Peripheral blood cell counts in Tie2-cre recipient mice (white) and Tie2-cre^+^FF1^+^ recipient mice (black) 16wk after transplantation. (n=5 mice in each group) (**C**) Marrow CD150^+^CD48^-^ HSPC frequencies in Tie2-cre recipient mice (left) and Tie2-cre^+^FF1^+^ recipient mice (right). (n=5 mice in each group) (**D**) Scheme of competitive marrow transplantation experiments. (**E**) Peripheral blood cell counts in recipient mice of Tie2-cre (CD45.2) and wild-type (CD45.1) donors (grey), and recipients of Tie2-cre^+^FF1 ^+^ (CD45.2) and wild-type (CD45.1) donors (black) 40wk after transplantation. (n=6 mice in each group) WBC: white blood cell; Hb: hemoglobin; Plt: platelet. * *P* <0.05

To test the possibility that mutant HSPCs may require a longer period of time to develop the disease phenotype in wild-type environment, we transplanted Tie2-cre^+^FF1^+^ or Tie2-cre marrow cells (CD45.2) together with wild-type competitor marrow cells (CD45.1) into lethally irradiated wild-type recipients and followed these mice up to 40 wks post transplantation. (Figure 3D) We did not observe any difference in peripheral blood cell counts between recipients of Tie2-cre donors and recipients of Tie2-cre^+^FF1^+^ donors in this competitive transplantation environment. (Figure 3E) These observations indicate that when both wild-type and JAK2V617F mutant marrow cells were transplanted together, mutant HSPCs by themselves are insufficient to develop a MPN phenotype, and a diseased microenvironment (e.g. with JAK2V617F-bearing vascular ECs) can overcome the competition between wild-type and JAK2V617F mutant cells and lead to the development of an overt MPN.

### The effects of co-existing wild-type cells on JAK2V617F mutant stem/progenitor cell function

With the establishment of these two cell competition models in either wild-type niche (with co-existing wild-type and JAK2V617F mutant cells but no MPN, as in Fig 2) or mutant niche (with mutant clonal expansion and overt MPN, as in Fig 3), we explored the competitive interactions between wild-type and mutant cells, and the impact of microenvironmental factors in driving JAK2V617F mutant clonal dominance in the development of MPNs.

First, we measured wild-type and JAK2V617F mutant cell proliferation *in vivo* by BrdU labeling (Figure 4A). Using Lin^-^cKit^+^Sca1^+^CD150^+^CD48^-^ as the more primitive HSC markers, we found that JAK2V617F mutant HSCs proliferated more rapidly than wild-type HSCs when transplanted separately into lethally irradiated wild-type recipients; however, when transplanted together, wild-type HSCs displayed a significantly higher proliferation rate than mutant HSCs. These results echoed that found *in vitro* (Figure 1). In contrast, there was no difference in cell proliferation between co-existing wild-type and JAK2V617F mutant Lin^-^cKit^+^Sca1^+^ (LSK) cells, and JAK2V617F mutant unfractionated marrow cells proliferated more rapidly than wild-type cells (Figure 4B-C). In addition, while there were equal numbers of wild-type and JAK2V617F mutant HSCs in recipients of 50% mutant and 50% wild-type donor cells, there were significantly more total mutant marrow cells than wild-type marrow cells. (Figure 4D) These findings suggest that competition between wild-type and JAK2V617F mutant cells varies at differing stages of differentiation, especially prominent at the most primitive cell level.

**Figure 4.**
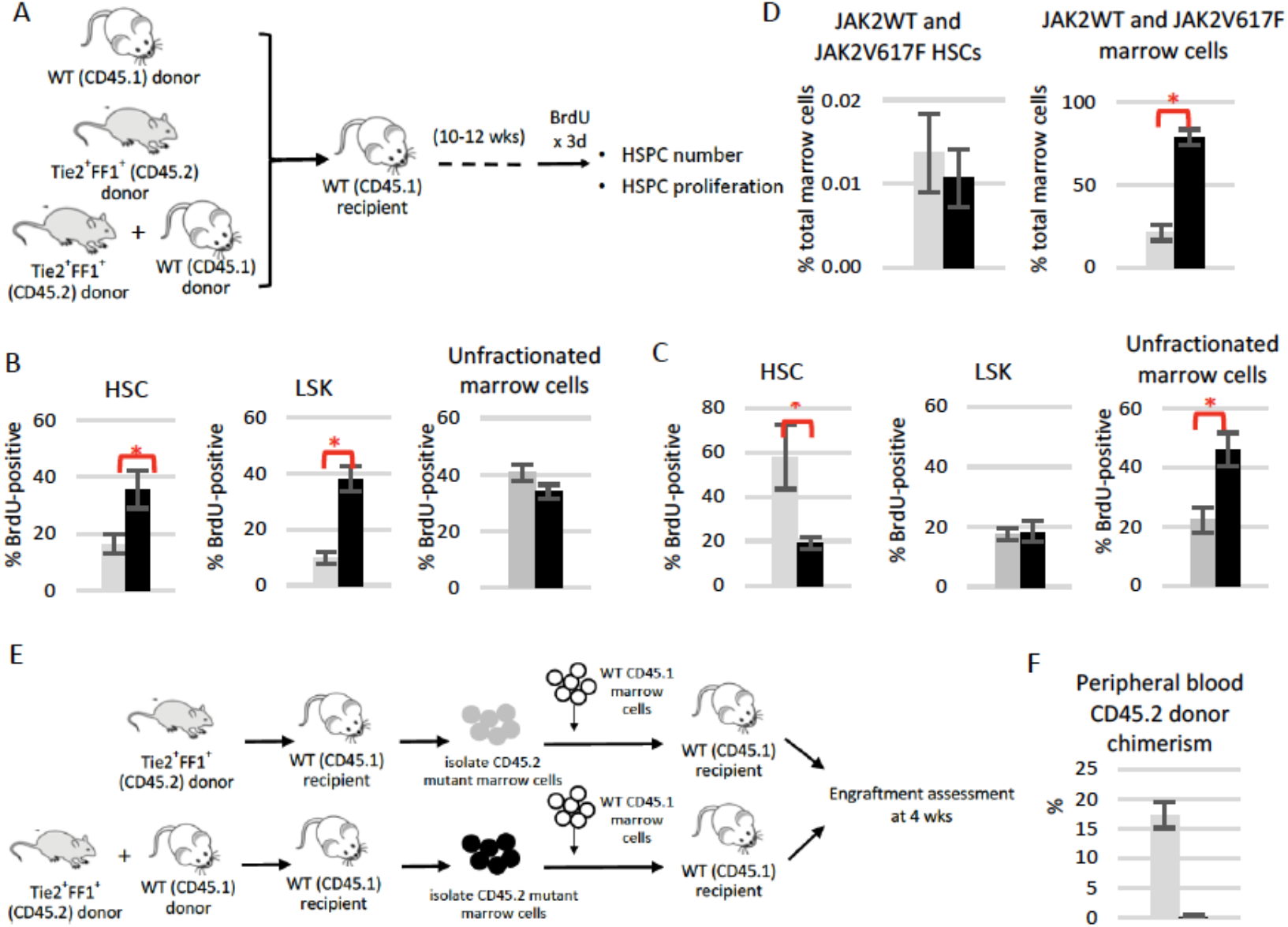
The presence of wild-type cells alters the behavior of JAK2V617F-mutant cells *in vivo.* (**A**) Scheme of marrow transplantation experiments. (**B-C**) Cell proliferation of wild-type (grey) and JAK2V617F mutant (black) HSCs (Lin^-^cKit^+^Sca1^+^CD150^+^CD48^-^), LSKs (Lin^-^cKit^+^Sca1^+^), and unfractionated whole marrow cells in direct transplantation (B) and in competitive transplantation (C). (n= 4-5 mice in each group) (**D**) Wild-type and JAK2V617F-mutant marrow HSCs (left) and unfractionated marrow cells (right) in recipients of both wild-type and mutant donor cells. (n=5 mice) (**E**) Scheme of marrow transplantation experiments. (**F**) Peripheral blood donor chimerism following competitive repopulation assay in which 5×10^5^ JAK2V617F marrow cells with (black) or without (grey) exposure to wild-type cell competition were injected together with 5×10^5^ competitor CD45.1 wild-type marrow cells into lethally irradiated CD45.1 wild-type recipients (n=4-5 in each group). * *P*<0.05

To evaluate the long-term consequences of such competitive interactions between wild-type and JAK2V617F mutant cells on mutant stem cell function, we performed a competitive repopulation assay in which we compared the engraftment potential of the JAK2V617F mutant marrow cells without the exposure to cell competition, to those with exposure to cell competition. (Figure 4E) At 4-week post-transplant, there was significantly more CD45.2 (JAK2V617F mutant) donor chimerism in the recipients of JAK2V617F mutant marrow donor without cell competition than in the recipients of JAK2V617F mutant marrow donor with cell competition. (Figure 4F) These data suggest that competition with wild-type cells suppressed JAK2V617F mutant stem cell function.

### A mutant microenvironment alters the competition between wild-type and JAK2V617F mutant cells

Next, we determined how microenvironmental signals influence the competition between wild-type cells and JAK2V617F mutant cells using the two cell competition models we have established (Figure 5A).

**Figure 5.**
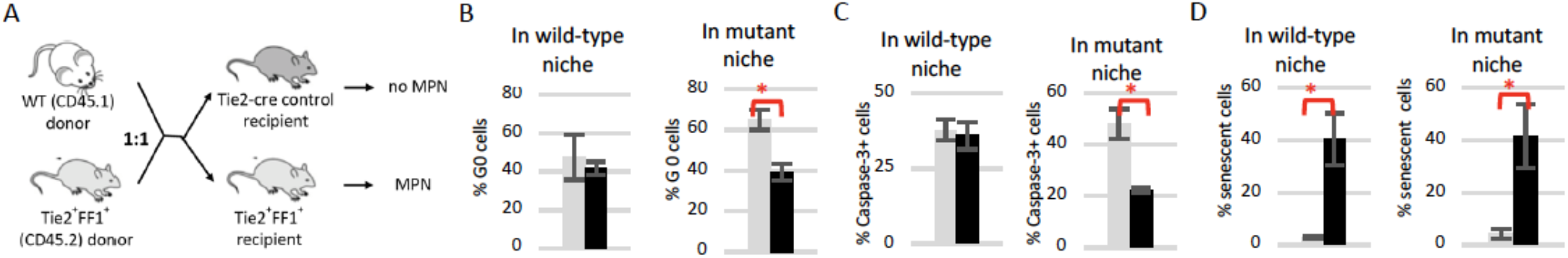
The microenvironment affects wild-type and JAK2V617F mutant cell competition. (**A**) Scheme of competitive marrow transplantation experiments where both wild-type and JAK2V617F mutant marrow donors were injected together into lethally irradiated Tie2-cre control mice (with wild-type vascular niche but no MPN) or Tie2-cre^+^FF1 ^+^ mice (with JAK2V617F mutant vascular niche and overt MPN). (**B-D**) G0 cell cycle status (B), cell apoptosis (C), and senescence (D) of wild-type (grey) and JAK2V617F mutant (black) CD150^+^CD48^-^ HSPCs in wild-type (left) and mutant niche (right). (n=4-6 mice in each group) * *P*<0.05

We examined cell cycle status of both mutant and wild-type CD150^+^CD48^-^ HSPCs, a highly enriched stem/progenitor cell population of which~ 20% display long-term repopulating capacity,^35^ using Hoechst33342 and Pyronin Y staining. We found that, consistent with the myeloproliferative phenotype we observed in the Tie2-cre^+^FF1^+^ recipients, JAK2V617F mutant cells were less quiescent (more cycling) than wild-type cells in the Tie2-cre^+^FF1^+^ recipients (with mutant ECs), while there were no differences in cell cycle status between wild-type and mutant HSPCs co-existing in control recipients (with normal ECs). (Figure 5B)

We then measured wild-type and JAK2V617F mutant cell apoptosis by activated-caspase-3 staining. While we did not detect any difference in the apoptosis levels between wild-type and JAK2V617F mutant HSPCs in the wild-type niche, mutant HSPCs had significantly less apoptosis compared to wild-type HSPCs in the mutant niche. (Figure 5C) These findings are consistent with our previous report that the mutant JAK2V617F-bearing vascular niche protects HSPCs from radiation-induced apoptosis.^34^

We also measured wild-type and JAK2V617F mutant cell aging by measuring senescence associated β-galactosidase (SA-β-Gal) activity using flow cytometry analysis. JAK2V617F mutant HSPCs demonstrated significantly higher senescence rates compared to wild-type HSPCs, in both wild-type niche and mutant niche. (Figure 5D)

### Immune responses triggered by cell competition between wild-type and JAK2V617F mutant cells

To determine the underlying molecular pathways that were differentially activated by wild-type and JAK2V617F mutant HSPC competition, we performed gene expression profiling on wild-type and JAK2V617F mutant Lin^-^ cKit^+^ HSPCs isolated from recipient mice in Figure 4A using RNA sequencing (RNA-seq). Principal component analysis and unsupervised hierarchical clustering (Figure 6A-B) revealed that mutant HSPCs with cell competition (i.e. transplanted together with wild-type cells) are distinct from mutant HSPCs without competition (i.e. transplanted alone), suggesting that the co-existing wild-type cells altered the transcriptomic profiling of JAK2V617F mutant HSPCs. 709 genes were differentially expressed (497 down-and 212 up-regulated) in JAK2V617F mutant HSPCs with cell competition compared to JAK2V617F mutant HSPCs without competition. Dysregulated immune response pathways are highly enriched in mutant HSPCs with cell competition compared to mutant HSPCs without cell competition. (Figure 6C)

**Figure 6.**
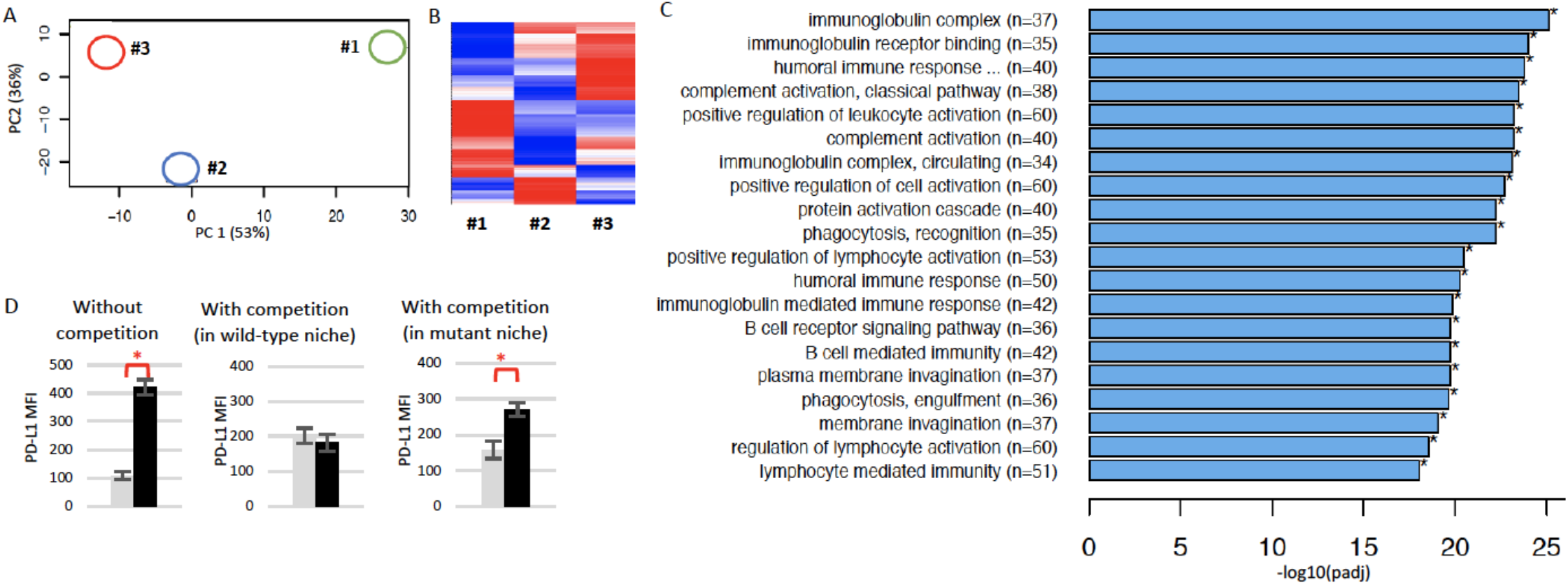
Immune response in wild-type and JAK2V617F-mutant cell competition. (**A-B**) Principal Component (PC) analysis (A) and heatmap of expression (B) of all differentially expressed genes (with an adjusted *P*-value<0.05) identified by RNA-seq data from *de novo* isolated wild-type Lin^-^cKit^+^ HSPCs transplanted alone (#1) (from 3 pooled mice), JAK2V617F mutant HSPCs transplanted together with wild-type cells (#2) (from 3 pooled mice), and JAK2V617F mutant HSPCs transplanted alone (#3) (from 2 pooled mice). (**C**) Top 20 differentially enriched GO terms in JAK2V617F mutant HSPCs transplanted together with wild-type cells (#2) compared to JAK2V617F mutant HSPCs transplanted alone (#3). *P* values are plotted as the negative of their logarithm. (**D**) Mean fluorescence intensity of PD-L1 staining on wild-type (grey) and JAK2V617F mutant (black) LSK cells in primary Tie2-cre^+^ control and Tie2-cre^+^FF1^+^ mice (left), when transplanted together into a wild-type niche (middle), and when transplanted together into a mutant JAK2V167F-bearing vascular niche (right). * *P*<0.05

Program death receptor-1 (PD-1) and one of its ligands, program death ligand 1 (PD-L1), constitute a major cellular tolerance mechanism. PD-L1 is expressed by many tumor cells. It binds to PD-1 expressed on activated T cells and results in T cell exhaustion, thereby blunting the anti-tumor immune response and promoting tumor growth.^42^ Consistent with a recent report that the oncogenic JAK2 activity induces PD-L1 expression,^43^ we found that PD-L1 surface expression was significantly increased on JAK2V617F mutant LSK cells from primary Tie2-cre^+^FF1^+^ mice compared to wild-type LSKs from Tie2-cre^+^ control mice. However, when transplanted together into a wild-type niche, there was no difference in PD-L1 expression between wild-type and mutant LSK cells; in contrast, when transplanted into a mutant niche (i.e. JAK2V617F-bearing vascular niche), PD-L1 expression was increased on JAK2V617F mutant LSK cells. (Figure 6D) Taken together, these data suggest that competition between wild-type and JAK2V617F mutant cells triggers a significant immune response, and deregulated PD-1/PD-L1 signaling may contribute to the stable co-existence of wild-type and mutant cells in a normal niche, and mutant clonal expansion in a mutant niche.

## Discussion

The co-existence of wild-type and JAK2V617F mutant cells in conditions that range from CHIP to chronic MPNs, which over long periods of time can remain stable or rapidly progress to frank malignancy, coupled with the high risk of relapse following curative allogeneic stem cell transplantation,^8–19^ all make MPN a unique model disease to study the early stages of clonal competition and tumor progression. Using both *in vitro* co-culture and *in vivo* competitive marrow transplantation assays, we demonstrate that the presence of wild-type cells alters the behavior of co-existing JAK2V617F mutant cells (Figure 1-2). While this appears to be in conflict with other reports that the JAK2V617F-positive MPN phenotype is transplantable and usually develops as early as 4 weeks following transplantation, in every such study 100% JAK2V617F-positive marrow cells were transplanted into wild-type recipients.^36–40^ These observations indicate that the “dosage” of mutant cells play an important role in the initiation of a MPN. It also suggests that competition between normal and neoplastic cell populations might be a zero-sum game in a wild-type microenvironment in which clonal expansion is limited by the availability of a microenvironment of finite size.

In contrast to the traditional view of cell competition in which more-fit cells out-compete their less-fit neighbors (“survival of the fittest”), our work suggests that competition between wild-type cells and JAK2V617F mutant cells may lead to their stable co-existence (“survival of the stable”). When 50% mutant marrow cells and 50% wild-type marrow cells were transplanted together into lethally irradiated wild-type recipient mice, the JAK2V617F mutant donor cells displayed engraftment similar to the wild-type donor cells, and the recipient mice had normal blood cell counts during up to ~10mo of follow up (Figure 2 and 3D-E). Quantitative measurement of cell proliferation *in vivo* revealed that, while mutant HSC proliferation was significantly decreased compared to the co-existing wild-type cells, there was no difference in cell proliferation between more differentiated LSK cells, and mutant unfractionated marrow cells proliferated more rapidly than wild-type cells. (Figure 4B-C) These data indicate that the co-existing wild-type cells can suppress JAK2V617F mutant stem cell proliferation while leaving more differentiated mutant cell expansion intact, leading to the stable co-existence of both wild-type and mutant cells without developing an overt MPN phenotype. Additional mutational or non-mutational processes (e.g. aging, inflammation) are likely required to enhance the competitiveness of JAK2V617F mutant cells to promote their clonal expansion.

In humans, cells carrying oncogenic mutations that are phenotypically silent for many years are not uncommon. It has long been recognized that the microenvironment surrounding tumor cells can provide tumor-suppressive signals to prevent mutant cells from expanding.^44,45^ Although the etiology of dysregulated hematopoiesis has been mainly attributed to the molecular alterations within the HSC compartment, abnormalities of the marrow niche are beginning to be recognized as an important factor in MPN development.^46–49^ To test whether a diseased microenvironment is such an “additional” factor required to enhance the competitiveness of JAK2V617F mutant cells, we used the Tie2-cre^+^FF1^+^ murine model which expresses the JAK2V617F mutation specifically in all hematopoietic cells and vascular ECs, so as to model the human diseases in which both the hematopoietic stem cells and ECs harbor the mutation.^22–27^ We found that, in contrast to the stable co-existence of both wild-type and mutant cells in a wild-type niche (Figure 2 and 3D-E), a JAK2 mutation-bearing niche can alter the interactions between wild-type and mutant cells to promote mutant clonal expansion and the development of a MPN (Figure 3A-C and Figure 5). These results indicate that cell competition in MPNs is a context-dependent process.

Patients with MPNs are characterized by a significant dysregulation of the immune system,^50,51^ and interferon-alpha, a potent immunostimulatory drug, can induce a considerable decrease in the mutant JAK2 allele burden in these patients.^52^ Recent work demonstrated that the JAK2V617F mutation increases PD-L1 expression, which can be used by the neoplastic cells to evade an antitumor immune response.^43^ In this study, we demonstrated that dysregulated immune response pathways are highly enriched in mutant HSPCs with co-existing wild-type cells (i.e. with competition) compared to mutant HSPCs alone (i.e. without competition) (Figure 6C), and there is a dynamic PD-L1 deregulation in JAK2V617F mutant HSPCs affected by their interactions with the neighboring wild-type cells and the microenvironment (Figure 6D). Therefore, not only the oncogenic mutation can cooperate with mechanisms to allow immune escape, but also the normal cells can employ the same immune mechanism to prevent mutant cell clonal expansion. Further understanding of the molecular mechanisms controlling the competitive interactions between normal and neoplastic stem cells, and how these mechanisms break down during cancer progression and relapse hold great potential for advances in treating cancer.

## ACKNOWLEDGEMENTS

This research was supported by the National Heart, Lung, and Blood Institute grant NIH R01 HL134970 (H.Z.), VA Career Development Award BX001559 (HZ), and VA Merit Award BX003947 (H.Z.). We thank Dr. Kenneth Kaushansky (Stony Brook University, NY) for valuable scientific discussions and critical review of the manuscript.

## AUTHOR CONTRIBUTION

M.C. performed various *in vitro* and *in vivo* experiments of the project; H. Zhang performed/assisted various flow cytometry and marrow transplantation experiments; H. Zhan conceived the projects, analyzed the data, interpreted the results, and wrote the manuscript.

## REFERENCES

1. Moreno E. Is cell competition relevant to cancer? Nat Rev Cancer 2008;8:141–7.

2. Green DR. Cell competition: pirates on the tangled bank. Cell Stem Cell 2010;6:287–8.

3. Johnston LA. Socializing with MYC: cell competition in development and as a model for premalignant cancer. Cold Spring Harb Perspect Med 2014;4:a014274.

4. Zwaka TP. Status Anxiety among Pluripotent Stem Cells? Dev Cell 2017;42:555–6.

5. Martincorena I, Fowler JC, Wabik A, et al. Somatic mutant clones colonize the human esophagus with age. Science 2018;362:911–7.

6. Martincorena I, Roshan A, Gerstung M, et al. Tumor evolution. High burden and pervasive positive selection of somatic mutations in normal human skin. Science 2015;348:880–6.

7. James C, Mazurier F, Dupont S, et al. The hematopoietic stem cell compartment of JAK2V617F-positive myeloproliferative disorders is a reflection of disease heterogeneity. Blood 2008;112:2429–38.

8. Lambert JR, Gale RE, Linch DC. The production of JAK2 wild-type platelets is not downregulated in patients with JAK2 V617F mutant-positive essential thrombocythaemia. Br J Haematol 2009;145:128–30.

9. Stein BL, Williams DM, Wang NY, et al. Sex differences in the JAK2 V617F allele burden in chronic myeloproliferative disorders. Haematologica 2010;95:1090–7.

10. Gale RE, Allen AJ, Nash MJ, Linch DC. Long-term serial analysis of X-chromosome inactivation patterns and JAK2 V617F mutant levels in patients with essential thrombocythemia show that minor mutant-positive clones can remain stable for many years. Blood 2007;109:1241–3.

11. Kroger N, Holler E, Kobbe G, et al. Allogeneic stem cell transplantation after reduced-intensity conditioning in patients with myelofibrosis: a prospective, multicenter study of the Chronic Leukemia Working Party of the European Group for Blood and Marrow Transplantation. Blood 2009;114:5264–70.

12. Rondelli D, Goldberg JD, Isola L, et al. MPD-RC 101 prospective study of reduced-intensity allogeneic hematopoietic stem cell transplantation in patients with myelofibrosis. Blood 2014;124:1183–91.

13. Guardiola P, Anderson JE, Bandini G, et al. Allogeneic stem cell transplantation for agnogenic myeloid metaplasia: a European Group for Blood and Marrow Transplantation, Societe Francaise de Greffe de Moelle, Gruppo Italiano per il Trapianto del Midollo Osseo, and Fred Hutchinson Cancer Research Center Collaborative Study. Blood 1999;93:2831–8.

14. Kroger N. Current Challenges in Stem Cell Transplantation in Myelofibrosis. Curr Hematol Malig Rep 2015;10:344–50.

15. Xie M, Lu C, Wang J, et al. Age-related mutations associated with clonal hematopoietic expansion and malignancies. Nat Med 2014;20:1472–8.

16. Genovese G, Kahler AK, Handsaker RE, et al. Clonal hematopoiesis and blood-cancer risk inferred from blood DNA sequence. N Engl J Med 2014;371:2477–87.

17. Jaiswal S, Fontanillas P, Flannick J, et al. Age-related clonal hematopoiesis associated with adverse outcomes. N Engl J Med 2014;371:2488–98.

18. McKerrell T, Park N, Moreno T, et al. Leukemia-associated somatic mutations drive distinct patterns of age-related clonal hemopoiesis. Cell Rep 2015;10:1239–45.

19. Jaiswal S, Ebert BL. Clonal hematopoiesis in human aging and disease. Science 2019;366.

20. Tiedt R, Hao-Shen H, Sobas MA, et al. Ratio of mutant JAK2-V617F to wild-type Jak2 determines the MPD phenotypes in transgenic mice. Blood 2008;111:3931–40.

21. Constien R, Forde A, Liliensiek B, et al. Characterization of a novel EGFP reporter mouse to monitor Cre recombination as demonstrated by a Tie2 Cre mouse line. Genesis 2001;30:36–44.

22. Sozer S, Fiel MI, Schiano T, Xu M, Mascarenhas J, Hoffman R. The presence of JAK2V617F mutation in the liver endothelial cells of patients with Budd-Chiari syndrome. Blood 2009;113:5246–9.

23. Rosti V, Villani L, Riboni R, et al. Spleen endothelial cells from patients with myelofibrosis harbor the JAK2V617F mutation. Blood 2013;121:360–8.

24. Yoder MC, Mead LE, Prater D, et al. Redefining endothelial progenitor cells via clonal analysis and hematopoietic stem/progenitor cell principals. Blood 2007;109:1801–9.

25. Teofili L, Martini M, Iachininoto MG, et al. Endothelial progenitor cells are clonal and exhibit the JAK2(V617F) mutation in a subset of thrombotic patients with Ph-negative myeloproliferative neoplasms. Blood 2011;117:2700–7.

26. Piaggio G, Rosti V, Corselli M, et al. Endothelial colony-forming cells from patients with chronic myeloproliferative disorders lack the disease-specific molecular clonality marker. Blood 2009;114:3127–30.

27. Helman R, Pereira WO, Marti LC, et al. Granulocyte whole exome sequencing and endothelial JAK2V617F in patients with JAK2V617F positive Budd-Chiari Syndrome without myeloproliferative neoplasm. Br J Haematol 2018;180:443–5.

28. Kent DG, Copley MR, Benz C, et al. Prospective isolation and molecular characterization of hematopoietic stem cells with durable self-renewal potential. Blood 2009;113:6342–50.

29. Kent DG, Li J, Tanna H, et al. Self-renewal of single mouse hematopoietic stem cells is reduced by JAK2V617F without compromising progenitor cell expansion. PLoS Biol 2013;11:e1001576.

30. Zhang Y, Lin CHS, Kaushansky K, Zhan H. JAK2V617F Megakaryocytes Promote Hematopoietic Stem/Progenitor Cell Expansion in Mice Through Thrombopoietin/MPL Signaling. Stem Cells 2018;36:1676–84.

31. Shapiro HM. Flow cytometric estimation of DNA and RNA content in intact cells stained with Hoechst 33342 and pyronin Y. Cytometry 1981;2:143–50.

32. Zhan H, Lin CHS, Segal Y, Kaushansky K. The JAK2V617F-bearing vascular niche promotes clonal expansion in myeloproliferative neoplasms. Leukemia 2018;32:462–9.

33. Lin CH, Kaushansky K, Zhan H. JAK2V617F-mutant vascular niche contributes to JAK2V617F clonal expansion in myeloproliferative neoplasms. Blood Cells Mol Dis 2016;62:42–8.

34. Lin CHS, Zhang Y, Kaushansky K, Zhan H. JAK2V617F-bearing vascular niche enhances malignant hematopoietic regeneration following radiation injury. Haematologica 2018;103:1160–8.

35. Kiel MJ, Yilmaz OH, Iwashita T, Yilmaz OH, Terhorst C, Morrison SJ. SLAM family receptors distinguish hematopoietic stem and progenitor cells and reveal endothelial niches for stem cells. Cell 2005;121:1109–21.

36. Etheridge SL, Roh ME, Cosgrove ME, et al. JAK2V617F-positive endothelial cells contribute to clotting abnormalities in myeloproliferative neoplasms. Proc Natl Acad Sci U S A 2014;111:2295–300.

37. Wernig G, Mercher T, Okabe R, Levine RL, Lee BH, Gilliland DG. Expression of Jak2V617F causes a polycythemia vera-like disease with associated myelofibrosis in a murine bone marrow transplant model. Blood 2006;107:4274–81.

38. Mullally A, Lane SW, Ball B, et al. Physiological Jak2V617F expression causes a lethal myeloproliferative neoplasm with differential effects on hematopoietic stem and progenitor cells. Cancer Cell 2010;17:584–96.

39. Akada H, Yan D, Zou H, Fiering S, Hutchison RE, Mohi MG. Conditional expression of heterozygous or homozygous Jak2V617F from its endogenous promoter induces a polycythemia vera-like disease. Blood 2010;115:3589–97.

40. Li J, Spensberger D, Ahn JS, et al. JAK2 V617F impairs hematopoietic stem cell function in a conditional knock-in mouse model of JAK2 V617F-positive essential thrombocythemia. Blood 2010;116:1528–38.

41. Crane GM, Jeffery E, Morrison SJ. Adult haematopoietic stem cell niches. Nat Rev Immunol 2017.

42. Flies DB, Chen L. The new B7s: playing a pivotal role in tumor immunity. J Immunother 2007;30:251–60.

43. Prestipino A, Emhardt AJ, Aumann K, et al. Oncogenic JAK2(V617F) causes PD-L1 expression, mediating immune escape in myeloproliferative neoplasms. Sci Transl Med 2018;10.

44. Stoker MG, Shearer M, O’Neill C. Growth inhibition of polyoma-transformed cells by contact with static normal fibroblasts. J Cell Sci 1966;1:297–310.

45. Bissell MJ, Hines WC. Why don’t we get more cancer? A proposed role of the microenvironment in restraining cancer progression. Nat Med 2011;17:320–9.

46. Walkley CR, Olsen GH, Dworkin S, et al. A microenvironment-induced myeloproliferative syndrome caused by retinoic acid receptor gamma deficiency. Cell 2007;129:1097–110.

47. Schepers K, Pietras EM, Reynaud D, et al. Myeloproliferative neoplasia remodels the endosteal bone marrow niche into a self-reinforcing leukemic niche. Cell Stem Cell 2013;13:285–99.

48. Arranz L, Sanchez-Aguilera A, Martin-Perez D, et al. Neuropathy of haematopoietic stem cell niche is essential for myeloproliferative neoplasms. Nature 2014;512:78–81.

49. Mager LF, Riether C, Schurch CM, et al. IL-33 signaling contributes to the pathogenesis of myeloproliferative neoplasms. J Clin Invest 2015;125:2579–91.

50. Tefferi A, Vaidya R, Caramazza D, Finke C, Lasho T, Pardanani A. Circulating interleukin (IL)-8, IL-2R, IL-12, and IL-15 levels are independently prognostic in primary myelofibrosis: a comprehensive cytokine profiling study. J Clin Oncol 2011;29:1356–63.

51. Skov V, Riley CH, Thomassen M, et al. Whole blood transcriptional profiling reveals significant down-regulation of human leukocyte antigen class I and II genes in essential thrombocythemia, polycythemia vera and myelofibrosis. Leuk Lymphoma 2013;54:2269–73.

52. Silver RT, Vandris K, Wang YL, et al. JAK2(V617F) allele burden in polycythemia vera correlates with grade of myelofibrosis, but is not substantially affected by therapy. Leuk Res 2011;35:177–82.

